# Cross talk between tumor stroma and cancer cells plays a critical role in progressive enrichment of cancer stem cell phenotype in primary breast tumors

**DOI:** 10.1101/2020.06.15.152355

**Authors:** Ninjit Dhanota, Amanjit Bal, Gurpreet Singh, Sunil K Arora

## Abstract

In order to delineate the underlying molecular mechanisms responsible for intra tumoral enrichment of BCSCs in aggressive breast tumors, firstly we evaluated the frequency and characteristics of breast cancer stem cells (BCSCs) within the tumor mass as well as in pathologically normal adjacent tissues in primary breast carcinomas of various clinical and histological grades. Then, we evaluated the expression profiles of various genes in non-cancer stem cells from these tumors to delineate the role played by cellular niche in de novo origin and/or expansion of intra-tumoral cancer stem cells.

The study included primary tumor and adjacent normal breast tissue specimens from chemotherapy-naïve breast carcinoma patients. The BCSCs, identified as Lin^-^CD44^+^CD24^-^ and aldehyde dehydrogenase 1 A1 positive were enumerated. The frequency of intra-tumoral BCSCs was correlated with various clinicopathological parameters of breast cancer. The flow-cytometrically sorted stromal cells and cancer cells from treatment naïve primary breast tumors were processed for gene expression profiling using a custom designed PCR array of genes known to facilitate cancer cell proliferation and disease progression.

The frequency of BCSCs within the tumor mass as well as in the adjacent normal tissue correlated significantly with histopathological and molecular grades of tumors indicating a direct relationship of BCSC with aggressive behavior of breast cancer. A significantly higher number of BCSCs was also detected in metastatic LN group as compared to non-metastatic LN. Further, a significantly increased expression of the genes associated with growth factors, cytokines & matricellular proteins in tumors with high BCSCs content (> 5%; Hi-BCSCs tumors) as compared to Lo-BCSC tumors (with <5% intratumoral BCSC content) suggested the possible contribution of stromal cells and cancer cells in intra-tumoral expansion of CSCs. Similarly, a significant up-regulation of genes associated with hypoxia and angiogenesis in Hi-BCSCs tumors further supported the role of hypoxic environment. The expression levels of genes associated with epithelial to mesenchymal transition also followed a similar pattern. On the other hand, downregulated SNAI1 gene (generally upregulated in onset of EMT) in stromal cells of Hi-BCSCs tumors suggests a post EMT environment in Hi-BCSCs tumors.

The findings suggest that the molecular crosstalk between the non-BCSC niche cells and the cancer stem cells within the breast cancer microenvironment directly contribute to formation of biologically conducive conditions for expansion of cancer stem cells.

## Introduction

The progression of breast cancer is highly unpredictable and dependent upon various factors. Advancements in early diagnosis have largely declined the overall mortality rate of patients with breast cancer, yet the long-term survival rate with metastatic and recurrent tumors has not improved significantly over the past several decades (1–3). Accumulating evidence indicates a small population of drug-resistant tumor cells known as “Cancer Stem cells (CSCs)”, showing increased metastatic/tumorigenic and stemness potential could be one of the root causes of relapse (4, 5). Although the exact involvement of CSCs in recurrence and metastasis is not well established yet, there are many putative molecular markers/factors that are under surveillance for their possible association (6, 7). Additionally, the study of breast cancer onset and progression has been focused mainly on the epithelial components of the tumor, while the surrounding stroma has attracted very little attention. New evidences have now emerged suggesting key interactions between tumor cells and stromal cells playing significant role in establishment and progression of tumors. Resistance to anti-cancer chemotherapies commonly involves four key mechanisms; (i) the over-expression of drug transporters/efflux pump (ii) the manipulation of apoptosis and senescence pathways by cancer cells (iii) the mechanical or stochastic factors and (iv) the presence of CSCs. The pan-resistance can be due to the remnant cancer cells after initial therapy, which makes it far more aggressive and typically unresponsive to any treatment (6). On the other hand, many reports demonstrate the dynamicity of cells to transform across a spectrum of epithelial and mesenchymal states, as opposed to undergoing a one-way EMP transition (7–11), and the transformed cells have same phenotypic markers as of CSCs. Besides, the cancer associated fibroblasts, endothelial cells and immune cells have also been shown to be associated with secretion of various factors that render the cancer more aggressive, possibly through induction of morphogenetic process called Epithelial to Mesenchymal transition (EMT) (12). Mechanisms like EMT are well reported for inducing generation of CSCs/BCSCs (also known as induced cancer stem cells; iCSCs/iBCSCs) through different signaling pathways (13–15). Subsequently, iCSCs/iBCSCs show enhanced expression of genes related to invasion, migration, metastasis and chemo-resistance (16). The interaction between cancer cells and stromal cells could initiate complex signaling cascade, which may be helping in the enrichment of CSCs in the tumor (17).

The critical analysis of the role played by BCSCs in breast cancer metastasis is mainly conceptual and speculative with a paucity of data defining the role of BCSCs in aggressiveness and progression of breast cancer. We hypothesized that the aggressiveness of breast cancer can be directly related to the enrichment of BCSCs in the tumor microenvironment. Multiple primary lesions may be formed as a result of the distant spread of BCSCs or conversion of normal tissue cells to cancerous cells under the influence of diseased microenvironment. Considering the crucial role played by BCSCs in cancer metastasis, we further investigated the probability of metastatic process to be a ‘stem cell phenomena’. The BCSCs may not take part in tumor metastasis directly but might be helping to switch on a cascade of pathways involved in metastatic process by stimulating surrounding non stem cancer cells (NSCCs) with help of various stromal factors and finally converting them to BCSCs or BCSC like cells. Thus, it became important to assess the association/role of various factors released by tumor stroma and the associated mechanisms that could be helping in enrichment of CSCs in the breast cancer. On analysis of the gene-expression data we observed that multiple factors released by the stromal cells seem to significantly influence the intensity and frequency of signals responsible for expansion of BCSCs.

## Materials and Methods

### Ethical statement

The study was approved by Institutional Committee on Stem cell Research (IC-SCR) ref no. PGI-IC-SCRT-56-2013/4516 dated 04.11.2013 and Institutional Ethics Committee (IEC) ref. no. PGI/IEC/2014/2151 dated 06.01.2014, of Post Graduate Institute of Medical Education and Research (PGIMER), Chandigarh, India. An informed written consent was obtained from all the participating human subjects.

### Study Design

Female patients within the age group of 18-70 years, undergoing mastectomy/lumpectomy as part of surgical management of breast carcinoma were recruited in the study. Female patients exposed to chemotherapy and male breast carcinoma cases were excluded. Surgically resected tumor specimens from 100 cases of breast carcinoma were included in the study.

### Specimen Collection

Total mastectomy and axillary clearance (TMAC) and/or lumpectomy specimens containing primary breast tumor as well as adjacent normal tissue (taken from the farthest distant site of the primary tumor without tampering the resection limits of tissue) were collected. Specimens collected in DMEM medium supplemented with antibiotics and Fetal Bovine Serum (FBS) were transported aseptically to the Molecular Immunology Laboratory. Specimens from reduction mammoplasties and other cosmetic surgeries were collected as normal control.

For studying migration markers on BCSCs we collected tissue specimens at different tissue distances from tumor (T- at primary tumor site, T.A.1- at 3mm away from tumor, T.A.2- at 1cm away from tumor, T.A.3- at 2cm away from tumor, T.D.- at 4 cm away from tumor margins) from 17 TMAC cases.

For flowcytometry (FCM) experiments, fresh specimens were processed for single cell suspensions and for Immunohistochemistry (IHC), paraffin blocks were prepared from formalin-fixed specimens from primary tumor as well as adjacent normal tissue. Serial sectioning of specimens was done for preparing paraffin blocks of adjoining normal tissue.

### Identification of BCSCs and sorting of cancer cells and stromal cells

Cells were stained with monoclonal antibody conjugates: Lin-FITC, CD44 - PE and CD24 -APC-H7 fluorochromes (BD Biosciences, USA), for identification of cancer cells, stromal cells and BCSCs by flow-cytometry (Supplementary Materials and methods). Cell populations based on the phenotypic markers: BCSCs as Lin^-^CD44^+^CD24^-^ cells, breast cancer cells as Lin^-^CD44^+^CD24^+^ & Lin^-^CD44^-^CD24^+^ and stromal cells identified as Lin^-^CD44^-^CD24^-^ & Lin^+^CD44^-^CD24^-^ were analyzed using FACS Diva software (BD Biosciences, USA) and sorted in a flow cytometer sorter (FACS Aria II, BD USA).

Additionally, single cell suspensions obtained from tissue specimens at different tissue intervals from tumor were stained with Lin-FITC, CD44-PE and CD24-APC H7, CXCR4-APC at 37°C for 15 minutes in a water bath. Cells were acquired on flow cytometer (FACS Aria II, BD USA) and populations were analyzed using FACS Diva software (BD Biosciences). BCSCs showing CXCR4 expression were gated and analyzed.

### Quantitative Real time PCR for gene expression and pathway analysis

The tumor specimens were categorized into two groups irrespective of histopathological grading: Hi-BCSCs tumors (with >5% of BCSCs) and Lo-BCSCs tumors (with <5% of BCSCs) for gene expression profiling of cancer cells (CC) and stromal cells (SC) sorted from these tumor tissue specimens. The differential expression profile of various genes for stromal factors in sorted populations of cancer cells and stromal cells from the tumor and adjacent tissues in two categories (Hi-BCSC & Lo-BCSC) were evaluated using a custom-designed PCR array (Supplementary Material). The PCR-array included 44 number of genes related to hypoxia, EMT, growth factors, cytokines and stromal factors, selected based on their roles in various pathways leading to expansion/origin of cancer stem cells (Supplementary Table 1-3). The protein-protein interactions and subsequent biological pathways affected by the differentially expressed genes among various study groups were analyzed by using KEGG (http://www.genome.jp/kegg/pathway.html); Reactome (http://www.reactome.org/) and String 9.1 (http://string-db.org/) databases.

### ALDH1A1 expression and scoring

The paraffin sections from all tumors and adjacent normal tissues were stained for ALDHA1 expression by immunohistochemistry (IHC). All staining runs were accompanied by appropriate control slides (normal human liver sections). ALDH1A1 staining was also performed on non-metastatic/ metastatic lymph node sections of 12 breast carcinoma patients.

All the stained slides were scanned by a pathologist in a blinded manner. The ratio of positive to negative cellular profiles was estimated as a percentage of all tumor cells in a slide. The intensity of ALDH1A1 expression was scored in tumor cells only. Stromal positivity (Leucocytes, Macrophages, Adipocytes, Mesenchymal cells present in stroma) was considered negative. Liver sections were used as a positive control for validating the ALDH1A1 staining on tumor sections. A histological score was obtained by counting the positive tumor cells with a score ranging from 0 to 4+. In order to classify patients into ALDH1A1 (+) and ALDH1A1 (-) groups, ALDH1A1 (+) tumor sections were scored as 4+ (≥50% positive tumor cells), 3+ (≥ 10% - <50%), 2+ (≥5% - <10%), 1+ (1-5%) and 0 (Negative). For the analysis, all 1+, 2+, 3+ and 4+ were considered positive.

### Statistical Analysis

Discrete categorical data was represented in the form of either an absolute number/percentage or mean±SE. Continuous data assumed to be normally distributed was written either in the form of mean ± SD or in the form of median and interquartile range. Mann-Whitney U test was used for statistical analysis of skewed continuous variables. Independent t-test was applied to compare normally distributed data of two groups. Data of more than two groups were compared using one-way ANOVA. To evaluate the correlation between different variables, Spearman correlation test was applied. P value <0.05 was considered statistically significant. The analysis was done with the help of GraphPad Prism 5 Version 5.03.

## Results

The present study was conducted in a prospective manner that included treatment naïve cases of breast cancer in female patients (Table 1), who had undergone either total mastectomy with axillary clearance (TMAC) or lumpectomy with axillary clearance (LAC).

**Table 1:** Clinicopathological characteristics of study subjects (n=100) Clinical and histopathological characteristics of 100 treatment naïve breast carcinoma cases.

| Variable | Number of Patients |
| --- | --- |
| Total | 100 |
| Mean Age (range) | 51 (26-82 years) |
| Mean tumor size (range) | 3.094 (1-6 cm) |
| pT1 ($\leq 2$cm) | 26 (28.5%) |
| pT2 ($> 2$ to $< 5$cm) | 61 (67.0%) |
| pT3 ($\geq 5$cm) | 4 (4.4%) |
| Axillary Lymph Nodal Status |  |
| Positive (%) | 43 (46.2%) |
| Negative (%) | 48 (52.7%) |
| Pathological Grade (%) |  |
| Grade 1 | 09 (9.6%) |
| Grade 2 | 28 (30.1%) |
| Grade 3 | 56 (60.2%) |
| Breast Cancer Molecular Subtype (%) |  |
| ER<sup>+</sup>/PR<sup>+</sup> | 38 (51.3%) |
| Triple Positive | 06 (8.1%) |
| HER 2<sup>+</sup> | 09 (12.2%) |
| Triple Negative | 21 (28.3%) |
| Ki67 index: |  |
| $\geq 14\%$ | 17 (23.3%) |
| $< 14\%$ | 56 (76.7%) |
| Type of Surgery: |  |
| Mastectomy | 79 |
| Lumpectomy | 21 |

Females presenting with breast carcinoma who were advised mastectomy/lumpectomy, within the age group of 18-70 years, were included in present study at Post Graduate Institute of Medical Education and Research, Chandigarh, India.

### Increased frequency of BCSCs correlates with aggressive behavior of breast cancer

The frequency of Lin^-^CD44^+^CD24^-^ cells in tumor samples was expressed as a percentage of total tumor cells by flow cytometry. The mean percent values of Lin^-^CD44^+^CD24^-^ (BCSCs) cell population in the tumor specimens showed a significant positive correlation with histopathological grades (Figure 1A & 1B), although it did not relate well with their molecular categories (Figure 1C). Surprisingly the adjoining normal tissue to tumor also showed comparable numbers of BCSCs in molecularly aggressive breast carcinomas (TNBC, Her2^+^ cases), suggesting a cellular contribution of BCSCs in aggressive behavior of these cancer subtypes (Figure 1C). Fifty five out of seventy-five (73.3%) tissues showed positivity for ALDH1A1 (Figure 1D) by IHC. Stromal positivity was not taken into consideration. An increasing trend in ALDH1A1 positivity was seen in histopathological grades (9.1% in grade I, 23.6% in grade II and 67.2% in grade III tumors) though the difference did not reach a significant level (Figure 1E, Table 2). Although no statistically significant difference was noted between ALDH1A1 positive and ALDH1A1 negative categories when the tumors were further stratified among four molecular subgroups as Luminal A, Luminal B, TN and HER2+ (p=0.528) (Figure 1E), yet we found higher ALDH1A1 expression in molecularly most aggressive breast cancer consisted of 42.9% of ALDH1A1 positive group (21 out of 49) suggesting association of ALDH1A1 with aggressive breast carcinoma.

**Fig. 1:**
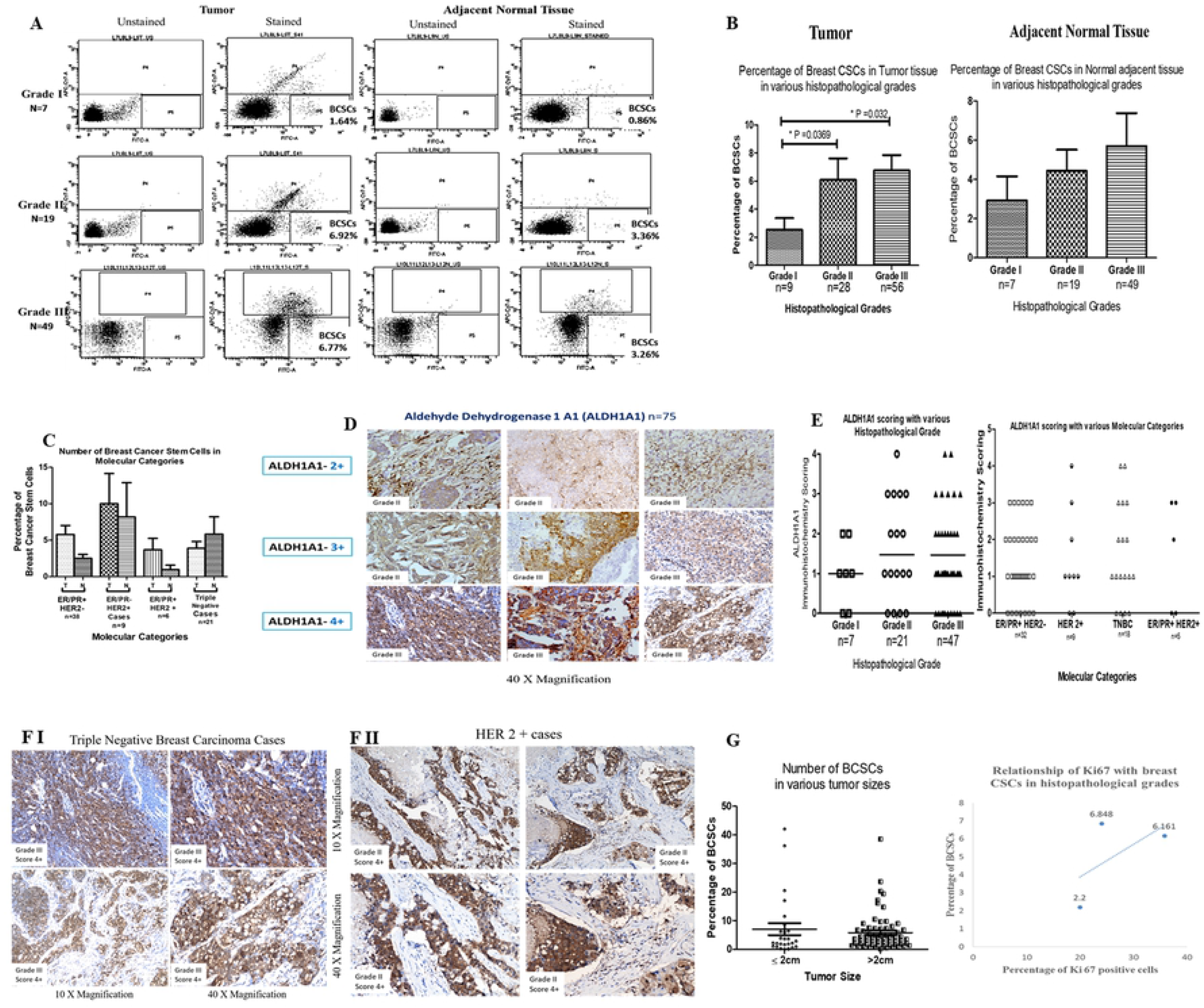
Increased percentage of BCSCs in tumor and adjacent normal breast tissues of primary breast carcinoma cases correlates with aggressive behavior of disease: (A). Representative flow cytograms showing comparative percentage of BCSCs in various histological grades in tumor and adjacent normal tissues. (B) Bar graphs showing comparative percentages (Mean±SEM) of BCSCs in tumor tissues indicating level of significance between Grade I vs Grade II (p=0.0369) and Grade I vs Grade III (p=0.032) tumors. No significant difference was found when percentage of BCSCs in adjacent normal tissue were compared. (C) Bar graphical representation of number of BCSCs (Mean ± SEM) in various molecular categories in tumor and adjacent normal tissues (ER/PR+ HER2-n=38; ER/PR-HER2+ n=9; ER/PR+ HER2+ n=6; ER/PR-HER2– n=21). (D) Representative photomicrographs showing scored immunohistochemical staining of tumor sections with ALDH1A1 antibody (40 X) in different histological grades. (E) Scatter plots showing comparison of ALDH1A1 IHC scores (Mean±SEM) in various histological grades of tumors (Grade I n=7; Grade II n=21; Grade III n=47) and molecular categories (ER/PR+ HER2-n=32; ER/PR-HER2+ n=9; ER/PR- HER2- n=18; ER/PR+ HER2 + n=5). (F) ALDH1A1 staining in molecularly aggressive breast carcinoma F-I. Triple Negative Breast Carcinoma category, F-II. Her 2 category. (G) Scatter plot showing comparative percentages of BCSCs in breast tumors of different sizes. No correlation found between percentages of BCSCs and tumor size (≤2cm and >2cm), however a positive trend line was observed when different histopathological grades were correlated with proliferation index.

**Table 2:** Comparison of clinico-pathological variables with ALDH1A1 positive or ALDH1A1 negative cases.

| Clinico-pathological parameters |  | ALDH1A1<sup>+</sup> CASES (n=26) | ALDH1A1<sup>-</sup> CASES (n=24) | p-value |
| --- | --- | --- | --- | --- |
| Mean age (range) ± SD |  | 52.5(26-82) ±13.025 | 53.5 (43-70) ± 8.937 | 0.796 |
| Mean tumor size ± SD |  | 3.32±1.291 | 2.46±1.117 | 0.042 |
| Tumor Grade | I | 5 (9.1%) | 2 (10%) | 0.371 |
|  | II | 13 (23.6%) | 8 (40%) |  |
|  | III | 37 (67.3%) | 10 (50%) |  |
| Molecular Sub groups | TP | 03 | 02 | 0.528 |
|  | TN | 14 | 04 |  |
|  | HER2<sup>+</sup> | 07 | 02 |  |
|  | HER2<sup>-</sup> | 25 | 7 |  |
| Metastatic Lymph Node | Present | 29 | 29 | 0.4035 |
|  | Absent | 11 | 7 |  |
| Median Lin<sup>-</sup>CD44<sup>+</sup>CD24<sup>-</sup> |  | 3.7 (1.40-9.15) | 4.2 (0.67-14.52) | 0.927 |

Clinicopathological characteristics were compared between ALDH1A1 positive and ALDH1A1 negative group. Significant difference between ALDH1A1 positive and ALDH1A1 negative groups was found in terms of tumor size.

We could not find any significant correlation in frequency of BCSCs and other clinicopathological parameters (e.g. Tumor size & Ki67) (Figure 1G, Table 2). The ALDH1A1 expression was associated with tumor size (p=0.042) but it did not correlate well with patient age, presence or absence of lymphovascular emboli and proliferation index (Table 2).

### BCSCs in metastatic lymph node and normal adjacent tissue indicate their role in invasiveness and metastasis

Lymph node (LN) metastasis is believably the initial step of distant metastasis and is directly related with a poor prognosis. A positive correlation between LN metastasis and number of BCSCs was observed (P = 0.0218 Spearmen Correlation r = 0.2120). The ALDH1A1 positivity was significantly higher in metastatic lymph node group (69%) compared to non-metastatic lymph node group (48%), although the difference was not statistically significant (Figure 2A). We found 30% lymph nodes with metastatic deposits showing ALDH1A1 positivity, however non metastatic LNs had no ALDH1A1 positive cells (N=3) (Figure 2B). Presence of BCSC like cells in normal adjacent tissue to tumors was also observed (Figure 2C & D), but no such cells were present in normal breast reduction mammoplasties. Cases presenting with high BCSC percentages by flow cytometry were shortlisted in grade II (n=4) and grade III (n=4) for ALDH1A1 staining by IHC. Out of these cases, two cases of grade II showed high ALDH1A1 expression (Supplementary Table 4, Figure 2C). Presence of BCSCs or BCSC like cells in adjacent areas of tumor suggests either infiltration or migration of these cells to normal areas or conversion of resident cells or cancer cells to these cells through various mechanisms. We also found expression of ALDH1A1 marker in histologically normal cells (n=3) adjacent to the tumor (Figure 2C), suggesting possible conversion of normal/cancer cells into BCSCs. Additionally, vimentin expression in 5-10% tumor cells suggests that possibly these cells could be undergoing EMT (Figure 2E). In order to evaluate the possible migration of BCSCs from tumor to adjacent normal tissue, we assessed the expression of CXCR4 on BCSCs. We observed percentage of BCSCs showing CXCR4 expression followed an increasing trend with the distance from the primary tumor site. The quantitative expression of CXCR4 (MFI) was found to be increasing as we move away from the primary tumor (Figure 2F, G).

**Fig. 2:**
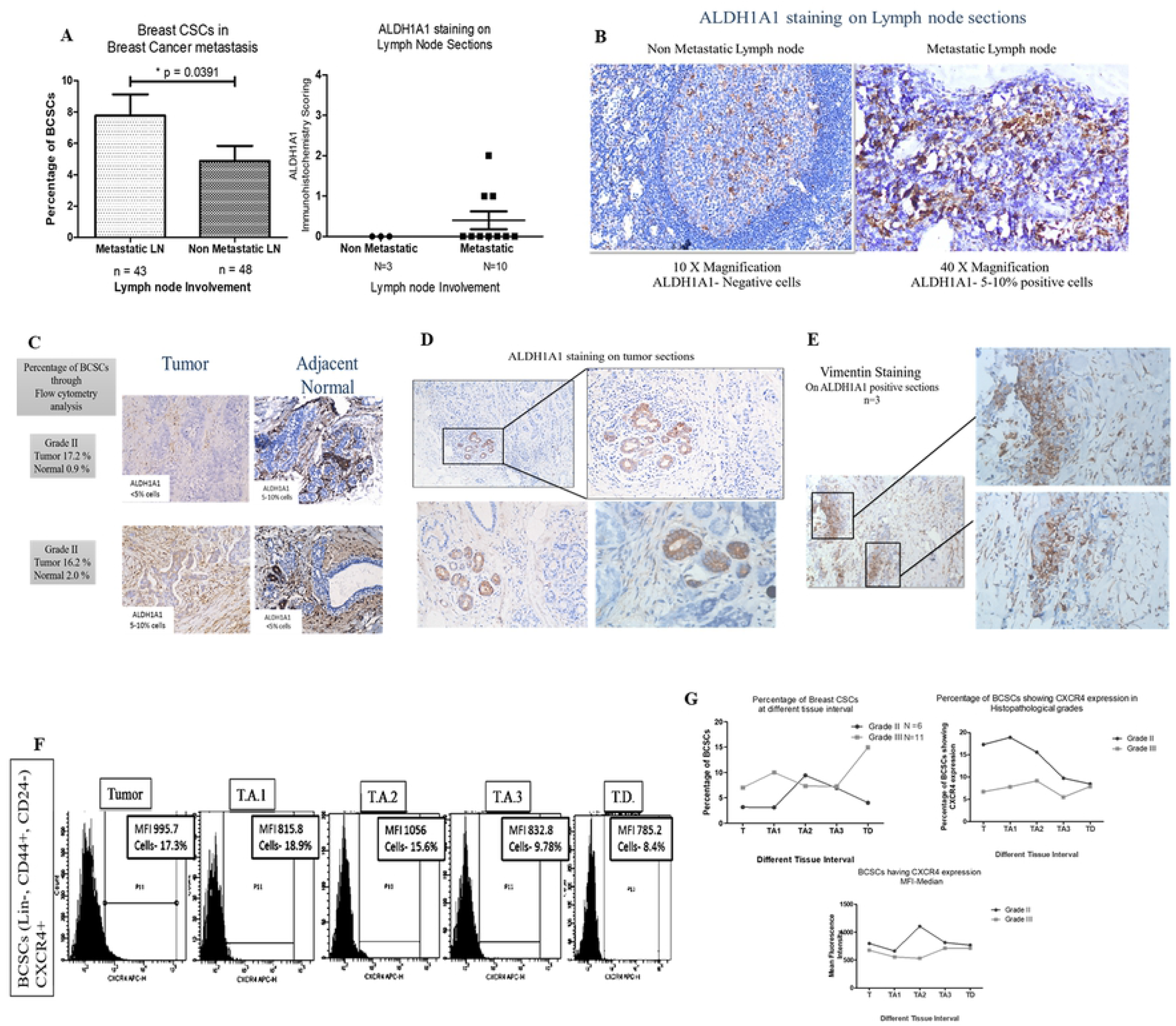
Status of BCSCs in breast cancer metastasis: (A) Bar graph representing number of BCSCs (Mean±SEM) in Metastatic lymphnodes (n=43) and non-metastatic lymphnodes (n=48) (flow cytometry) and a scatter plot showing ALDH1A1 staining scores on lymph node sections in metastatic (n=10) vs non-metastatic (n=3) nodes. (B) Microphotographs of ALDH1A1 staining in non-metastatic and metastatic lymph node sections. (C) Representative IHC staining of ALDH1A1 on shortlisted tumor cases showing high percentages of BCSCs in tumor and adjacent normal tissues by flow cytometry, indicative of field cancerization. (D) Presence of ALDH1A1+ cells within well-arranged histologically normal mammary ducts in tumor vicinity, IHC staining (E) Immunohistochemical staining for vimentin on tumor sections showed ALDH1A1+ normal cells in tumor vicinity indicating occurrence of EMT mechanism for conversion of NSCC to BCSCs. (F) Representative flow histograms showing percentage of BCSCs at different tissue levels in grade II and grade III tumors. (G) (upper left) Line plot representing mean distribution of BCSCs (Grade II and Grade III); (upper right) BCSCs showing CXCR4 expression at different tissue levels; (lower panel) CXCR4 MFI of BCSCs within Grade II and Grade III tumors at different tissue intervals from primary tumor tissue. (NSCC: Normal stem/cancer cells)

### Differential gene expression profiling in Cancer cells (CC) and stromal cells (SC) from Hi-BCSCs tumors (Test) and Lo-BCSCs tumors (Control)

Differential gene expression profile of cancer cells and stromal cells were analyzed in 20 samples of different histopathological grades and different BCSCs frequency. The mean values of BCSCs across Hi-BCSC and Lo-BCSC tumors were 16.68% and 2.52% respectively (Fig. 3A). Thirty-nine genes were found to be up-regulated in stromal cells and cancer cells out of which 26 genes showed two-fold up-regulation in Hi-BCSC tumors when compared to Lo-BCSC groups (Fig. 3B-G, Table 3). These included genes associated with hypoxia (HIF1A, ARNT, EPAS1, SIAH1, ZEB1, TAZ), inflammatory cytokines (IL-6, IL-8, TGF-β1, TNF-α), growth factors (VEGFA, FGF2, PDGFD, HGF), epithelial to mesenchymal transition (TWIST1, SOX9, CDH1, CDH2, VIM), and matricellular proteins (LUM, COL6A3, HAS2, POSTN, TNC, SPP1, SPARC).

**Fig. 3:**
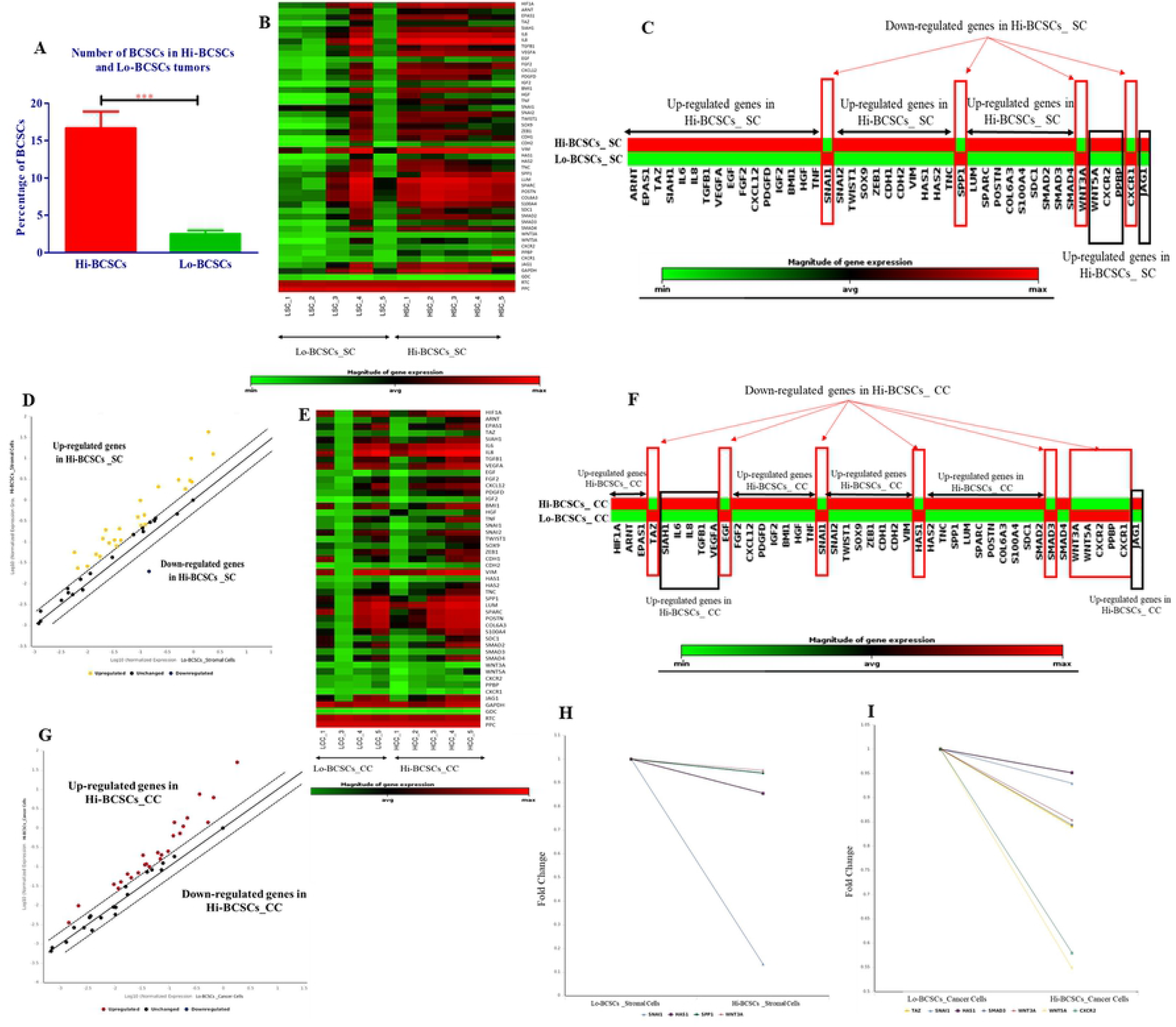
Differential gene expression in tumors with High BCSCs vs Low BCSCs: (A) Bar diagram showing percentage (Mean±SEM) of BCSCs in Hi-BCSC and Lo-BCSC breast tumors indicating significant difference in percentage of BCSCs in Hi-BCSCs (16.68%) vs Lo-BCSCs (2.52%) (p=0.0002) breast tumors. (B) Heat map showing the differential gene expression (fold change) in stromal cells of Hi-BCSCs_SC group (Test group) as compared to Lo-BCSCs_SC group (Control group). (C) Cluster diagram showing differential gene expression in stromal cells of Hi-BCSCs tumors (Test) vs. Lo-BCSCs tumors (Control). (D) Scatterplot showing the differential gene expression of up-regulated and down-regulated genes in stromal cells of Hi-BCSCs (Hi-BCSCs _SC) as compared to Lo-BCSCs (Lo-BCSCs_SC) tumors. (E) Heat map showing the differential gene expression (fold change) in cancer cells in Hi-BCSCs_CC group (Test group) as compared to Lo-BCSCs_CC group (Control group). (F) Cluster gram showing differential gene expression in cancer cells of Hi-BCSCs tumors (Test) vs. Lo-BCSCs tumors (Control). (G) Scatterplot showing differential gene expression of up-regulated and down-regulated genes in Hi-BCSCs_CC group as compared to Lo-BCSCs_CC group. (H) Line diagram showing down-regulation in genes (fold change) involved in EMT mechanism, Extracellular matrix protein contributing to aggressive behavior of disease and signaling molecules in stromal cells of Hi-BCSCs tumors as compared to Lo-BCSCs tumors. (I) Fold change in genes involved in EMT mechanism, chemokine receptors, Extracellular matrix protein contributing to aggressive behavior of disease and signaling molecules in cancer cells of Hi-BCSCs tumors as compared to Lo-BCSCs tumors. Low level of gene expression is represented by green color and high level of gene expression is represented by red color.

**Table 3A:**
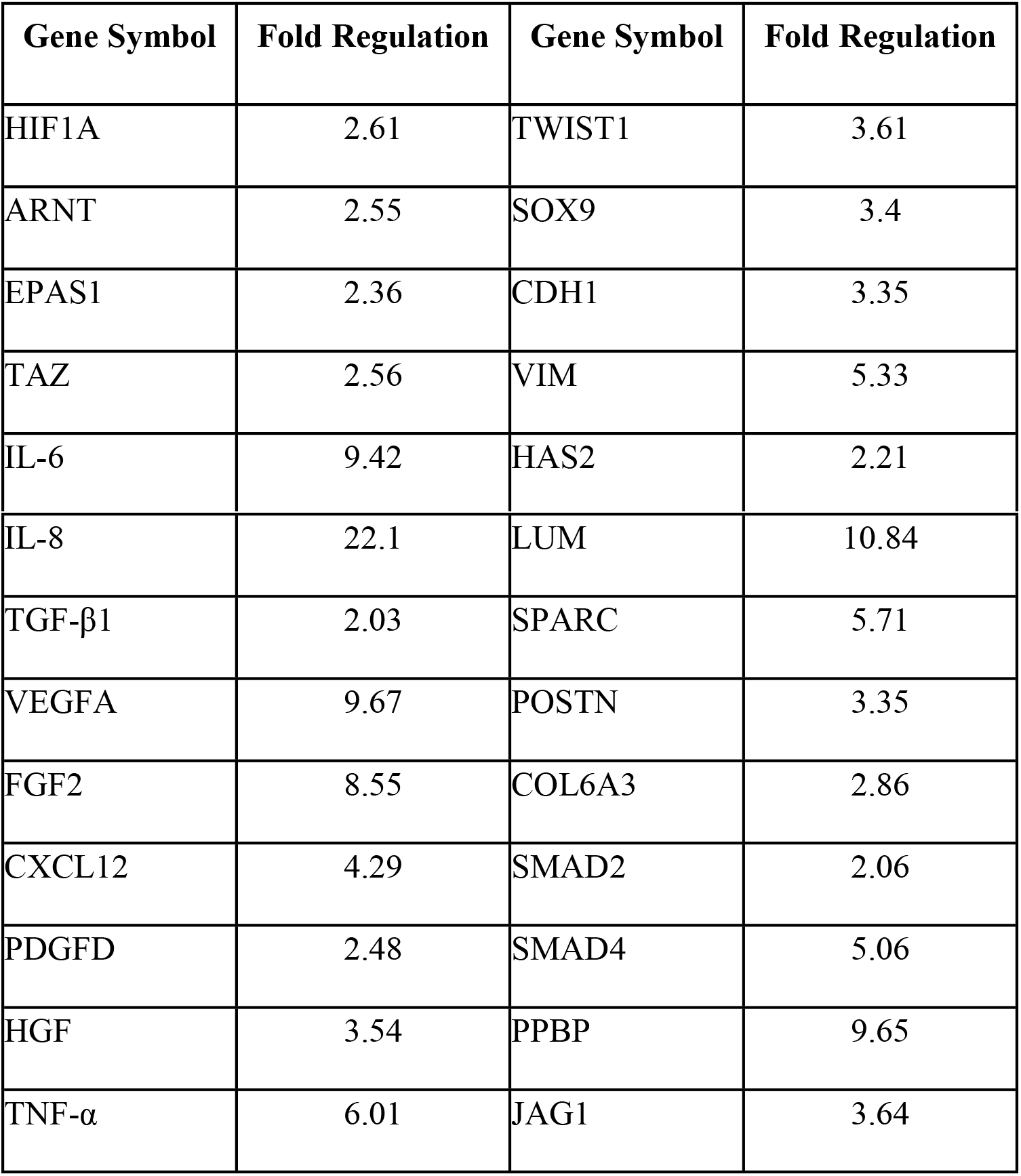
Gene symbols and their up-regulated expression (fold change) in stromal cells Hi-BCSCs_SC group (Test) as compared to Lo-BCSCs_SC group (Control).

| Gene Symbol | Fold Regulation | Gene Symbol | Fold Regulation |
| --- | --- | --- | --- |
| HIF1A | 2.61 | TWIST1 | 3.61 |
| ARNT | 2.55 | SOX9 | 3.4 |
| EPAS1 | 2.36 | CDH1 | 3.35 |
| TAZ | 2.56 | VIM | 5.33 |
| IL-6 | 9.42 | HAS2 | 2.21 |
| IL-8 | 22.1 | LUM | 10.84 |
| TGF-β1 | 2.03 | SPARC | 5.71 |
| VEGFA | 9.67 | POSTN | 3.35 |
| FGF2 | 8.55 | COL6A3 | 2.86 |
| CXCL12 | 4.29 | SMAD2 | 2.06 |
| PDGFD | 2.48 | SMAD4 | 5.06 |
| HGF | 3.54 | PPBP | 9.65 |
| TNF-α | 6.01 | JAG1 | 3.64 |

**Table 3B:** Gene symbols and their up-regulated expression (fold change) in cancer cells Hi-BCSCs_CC group (Test) as compared to Lo-BCSCs_CC group (Control).

| Gene Symbol | Fold Regulation | Gene Symbol | Fold Regulation |
| --- | --- | --- | --- |
| IL-8 | 27.72 | TWIST1 | 3.54 |
| IL-6 | 22.18 | ZEB1 | 3.08 |
| LUM | 20.36 | HAS2 | 3.06 |
| COL6A3 | 11.00 | SMAD4 | 3.00 |
| VIM | 9.16 | SIAH1 | 2.73 |
| POSTN | 8.28 | SPARC | 2.64 |
| HIF1A | 5.94 | BMI1 | 2.55 |
| CXCL12 | 5.83 | PDGFD | 2.51 |
| SPP1 | 5.20 | TNF-α | 2.50 |
| HGF | 4.49 | CDH2 | 2.44 |
| VEGFA | 4.45 | ARNT | 2.33 |
| FGF2 | 3.68 | TNC | 2.26 |
| S100A4 | 3.65 | SMAD2 | 2.25 |

Among the four down-regulated genes in stromal cells (Fig. 3H) only 1 gene was found to be significantly (≥2 fold) down-regulated in Hi-BCSC_SC group as compared to Lo-BCSC_SC group. The down-regulated genes included the genes involved in Epithelial to mesenchymal transition (SNAI1; seven-fold), genes of extracellular matrix proteins (HAS1, SPP1) involved in aggressive behavior of disease and a signaling molecule (WNT 3A). Similarly, in cancer cells downregulated genes are involved in Epithelial to mesenchymal transition (SNAI1; TAZ), Chemokine receptors (CXCR2), genes of extracellular matrix proteins (HAS1) involved in aggressive behavior of disease and signaling molecules (WNT 3A, WNT5A, SMAD 3).

Deep analysis and close scrutiny of gene expression data revealed that among the over-expressed genes in Hi-BCSC tumors, 19 genes were common as upregulated both in stromal cells as well as cancer cells, whereas 7 genes were exclusively over-expressed by either stromal cells (Figure 4A; Stromal cell’s box) or cancer cells (Fig. 4A; Cancer cell’s box). Only one gene, SNAI1 was found to be significantly down-regulated in stromal cell compartment of Hi-BCSC tumors as compared to Lo-BCSC tumors (Fig. 4B). In order to evaluate the role being played by various cellular components known to be part of BCSC niche, we mainly concentrated on role of hypoxia, ECM components, cytokines, chemokines and growth factors (Fig. 4B & E). The analysis revealed a significant positive correlation between percentage of intratumoral BCSCs with expression level of VEGFA (Spearman’s ρ= 0.552, p<0.05), CXCL12 (Spearman’s ρ= 0.453, p<0.05) and IL-6 (Spearman’s ρ=0.509, p<0.05) in the stromal cells (Fig. 4C, 4D, 4F) (Table 4). The correlation data is tabulated in Table 4.

**Fig. 4:**
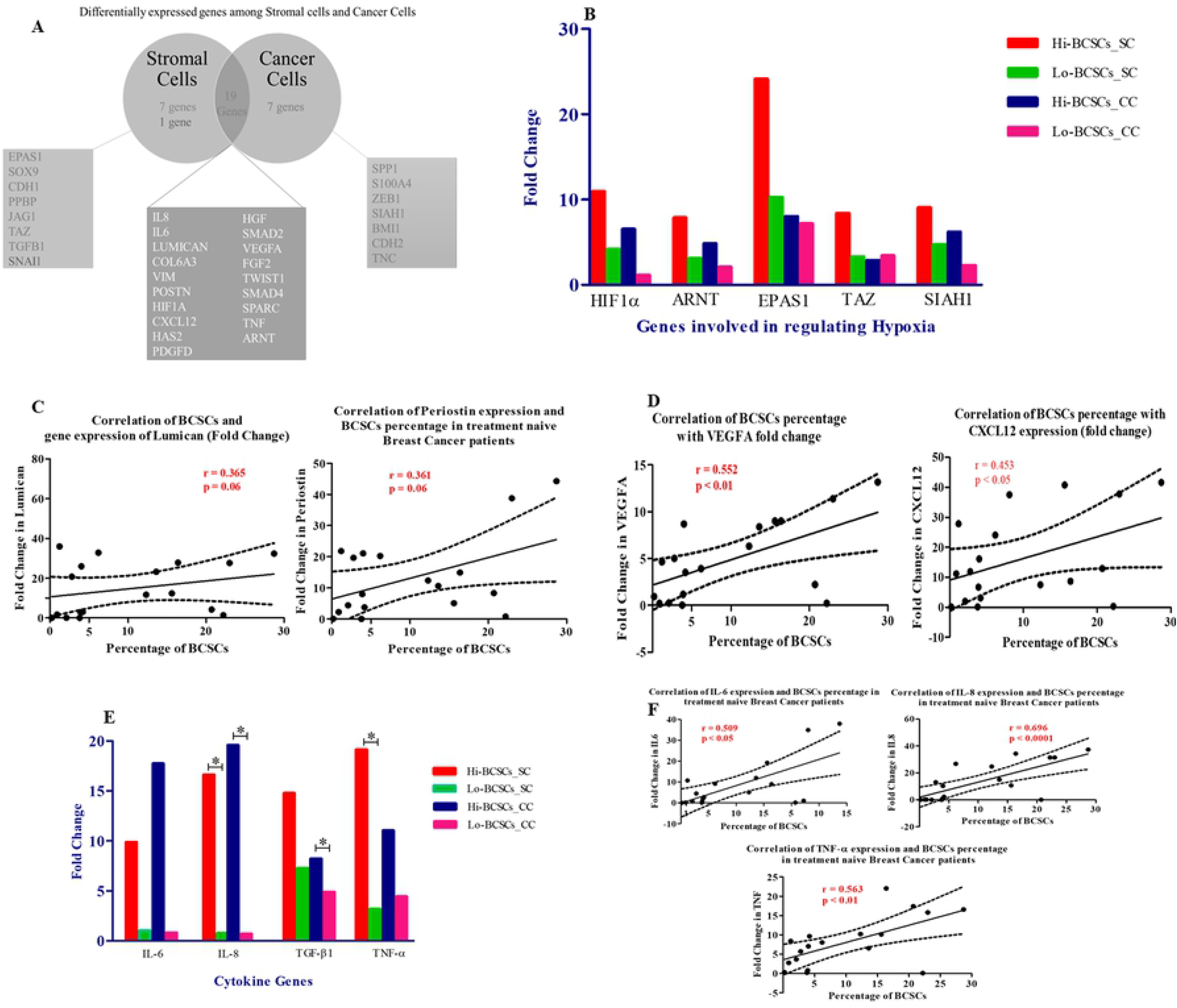
Differentially expressed genes among stromal cells and cancer cells of Hi-BCSCs and Lo-BCSCs tumors: (A). Flow diagram showing description of the differentially expressed genes among stromal cells and cancer cells sorted from Hi-BCSC tumors vs Lo-BCSC tumors. A total of 19 genes were found to be commonly over-expressed by cancer cells as well as stromal cells, whereas 7 genes were exclusively over-expressed by either stromal cells or cancer cells. Only one gene, SNAI1 was found to be significantly under-expressed in stromal cell compartment of Hi-BCSCs tumors as compared to Lo-BCSCs tumors. (B) Expression profiles of hypoxia related genes in Hi-BCSC tumors and Lo-BCSC tumors with respect to adjacent normal tissue (Control). Fold change at gene expression levels of HIF1α, ARNT, EPAS1, TAZ and SIAH1 in stromal cells and cancer cells of Hi-BCSCs tumors and Lo-BCSCs tumors as compared to adjacent normal tissue gene expression levels. (C) Correlation of BCSCs percentage with fold change in gene expression of Lumican and Periostin indicating status of ECM components in BCSC expansion. (D) Correlation of BCSCs percentage with fold change in gene expression of VEGFA and CXCL12 indicating role of chemokines and growth factors released by stromal cells and cancer cells in BCSC expansion. (E) Bar diagram showing fold change in gene expression levels of IL-6, IL-8, TNF-α and TGF-β1 in stromal cells and cancer cells of Hi-BCSCs tumors and Lo-BCSCs tumors as compared to adjacent normal tissue gene expression levels indicating association of inflammatory cytokines with BCSC expansion. (F) Correlation of BCSCs percentage with fold change in gene expression of IL-6, IL-8 and TNF-α

**Table 4:** Spearman’s rank correlation coefficients of differentially expressed genes (fold change) and BCSCs percentage.

|  | **Spearman's Rank Correlation Coefficient($\rho$)** | **p value** |
| --- | --- | --- |
| **HIF1α** | 0.279 | 0.124 |
| **ARNT** | 0.313 | 0.096 |
| **EPAS1** | 0.068 | 0.390 |
| **TAZ** | 0.014 | 0.477 |
| **SIAH1** | 0.111 | 0.326 |
| **LUM** | 0.365 | 0.062 |
| **COL6A3** | 0.279 | 0.124 |
| **POSTN** | 0.361 | 0.064 |
| **SPP1** | 0.135 | 0.291 |
| **HAS2** | 0.235 | 0.166 |
| **SPARC** | 0.239 | 0.163 |
| **TNC** | 0.292 | 0.112 |
| **HGF** | 0.207 | 0.395 |
| **VEGFA\*** | 0.552 | 0.014 |
| **FGF2** | 0.335 | 0.08 |
| **PDGFD** | 0.274 | 0.257 |
| **CXCL12\*** | 0.453 | 0.026 |
| **PPBP** | 0.100 | 0.342 |
| **IL-6\*** | 0.509 | 0.026 |

## Discussion

Failure of existing treatment modalities and presence of residual disease in cancer has been linked to presence of cancer stem cells in previous reports, with some studies reporting their role in resistance to conventional therapy while others supporting their role in relapse and metastasis (18). Our present study indicates a direct association of aggressive behavior of breast cancer with the frequency of CSCs in the tumor. Proposed explanations for the observed increase in number of CSCs in aggressive tumors could be attributed to many reasons viz. a) abnormal expression of factors associated with CSC proliferation in tumor, b) conversion of non-cancer cells to CSCs (de-differentiation) by aberrant/dysregulated signaling pathways, c) the intensified signals responsible for intra-tumoral expansion of BCSCs under the influence of stroma or positive feed-back by BCSCs themselves. Although the possible role of various intrinsic signals stemming from the tumor microenvironment on the expansion of BCSCs have been sparsely speculated with the help of some *in vitro* and animal studies, yet the behavior of diseased stroma at different grades of cancer in primary tumors in humans have not yet been fully elucidated (13, 19). The specific mechanisms by which cancer cells and surrounding non cancer cells influence the CSCs in terms of their expansion and formation of primary and metastatic niche sites are currently under investigation. Differentially expressed profile of genes revealed that down-regulated genes (SNAI1, Wnt ligands) in Hi-BCSC vs Lo-BCSC tumors were presenting post EMT condition, while the up-regulated genes were associated with hypoxia and inflammation along with concomitant overexpression of growth factors, suggesting a continuous surge of stimuli from stromal cells and cancer cells contributing to intra-tumoral expansion of CSCs.

Direct cell-cell interactions between the stromal cell compartment, cancer cells and CSCs, as well as signaling pathways mediated through the expression and secretion of a range of growth factors and cytokines play an important role in the maintenance of the CSC pool within the niche and in overall tumor growth (20, 21). As the tumor progresses, the normal stroma undergoes desmoplastic reaction through drastic changes and expansion (22). This desmoplastic expansion of stroma results in increased signals coming from activated fibroblasts, myofibroblasts and inflammatory cells, resulting in ECM remodeling and neovascularization (22). Observations from our present study support above mentioned notion, as we found a significantly increased expression of growth factors like VEGFA, FGF2, PDGFD, HGF that are secreted by stromal cells (fibroblasts, endothelial cells, myofibroblasts, immune cells) and cancer cells in Hi-BCSC tumors as compared to Lo-BCSC tumors. This suggests the critical role played by these growth factors in acquisition of CSC-like phenotype, thereby maintaining the CSC population within the tumor. The increased expression of S100A4 gene, which is associated with myofibroblasts, indicates that the stromal cells might help in inducing migratory properties in CSCs and thereby playing a significant role in driving metastasis. In addition, the elevated expression of CXCL12 by stromal cells in the vicinity of tumor along with upregulated expression of CXCR4 on cancer cells as well as CSCs (unpublished personal observation), complements the supportive role of the niche. Furthermore, various studies have shown the importance of endothelial cells in regulating the CSC numbers. In one of the previously reported studies, removal of endothelial cells from the CSC niche resulted in a decrease in the CSC numbers, suggesting dependence of CSCs on various cells of the niche cells for their maintenance (23, 24). We found an increased expression of BMI-1 and JAG-1 genes in stromal cells isolated from Hi-BCSC tumors, which supports the self-renewability and migratory potential of CSCs. These results corroborate the findings of earlier reports which indicated that the endothelial cells and immune cells secrete various factors like BMI1, JAG1, VEGFA and EGF helping in growth of cancer stem cells (25–28).

A number of experimental, clinical and epidemiological studies have revealed that the chronic inflammation contributes positively to cancer progression (28–30). Cancer-associated inflammation is mediated by infiltration of leucocytes and secretion of various cytokines such as TNF-α, IL-6, IL-8, TGF-β1, IL-10 etc. (31, 32). The inflammatory cytokines secreted by a broad range of cells, including immune cells, fibroblasts and endothelial cells present in CSC niche (31, 32) strongly support the association of cytokines with the maintenance and expansion of intra-tumoral CSC pool (33–35). A significantly increased expression of IL-6, IL-8, TGF-β1 and TNF-α in stromal cells and cancer cells isolated from primary breast tumors having high percentage of BCSCs in our study further support the significant role of inflammatory tumor environment in expansion of CSCs (Fig. 5A).

**Fig. 5:**
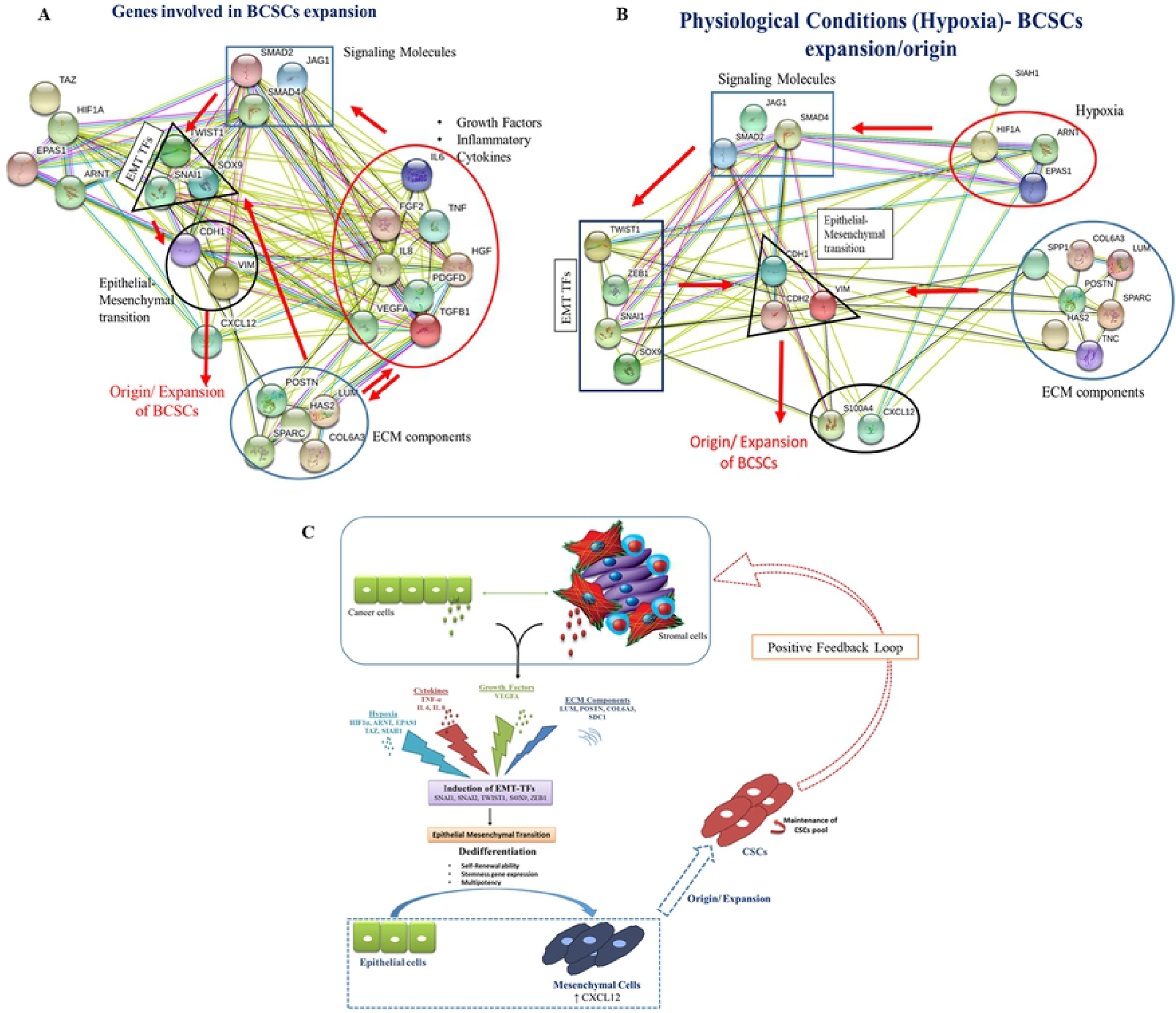
Proposed model of BCSCs origin/expansion. (A) String analysis showing gene interactions involved in BCSCs origin/expansion. Interactions of different growth factors, cytokines, ECM component genes with EMT regulatory genes, thereby helping in acquisition of BCSC phenotype. (B) String analysis suggesting role of hypoxia/hypoxia related genes in origin/expansion of BCSC phenotype. (C) Proposed mechanism suggesting cross talk between stromal cells and cancer cells initiate a cascade of events leading to increase in various intratumoral factors contributing to intra-tumoral expansion of CSCs. Based on our findings, it is proposed that the growth factors along with inflammatory cytokines through Notch and SMAD signaling induce EMT-transcription factors, which in turn initiate epithelial to mesenchymal transition. EMT might be one of the possible mechanisms responsible for acquisition of BCSCs phenotype.

A close link between epithelial to mesenchymal transition (EMT) and acquisition of CSC like properties, has enabled a greater understanding of the molecular mechanisms underlying the expansion and maintenance of CSCs in tumor masses (13, 19, 36). The stimuli that regulate EMT in cancer like specific signaling pathways (e.g. TGF-β1, Hedgehog, Wnt and Notch), cellular cross talk and hypoxic microenvironment, also include factors that are known to play key roles in the maintenance of CSCs. Besides the growth factors such as HGF, VEGFA, EGF, FGF2 and TGF-β1, the hypoxia related factors such as HIF1A, ARNT, EPAS1, TAZ, SIAH1 and neovascularization are also associated with EMT induction. Our gene-expression data revealed that the factors known to regulate the EMT transcription factors such as TWIST1, SOX9, SNAI1, SNAI2 and mesenchymal markers like Vimentin and N-cadherin, were all found to be significantly highly expressed in Hi-BCSC tumors as compared to L0-BCSC tumors, which suggests the significant role being played by EMT induction in expansion of CSCs in the breast cancer (Fig. 5B). This is in sync with the earlier reports supporting that induction of EMT helps in acquisition of CSC like properties during tumor progression (13, 14). Although SNAI1 gene expression was found to be significantly down-regulated in cancer cells as well as in stromal cells, yet it might suggest a post-EMT condition in Hi-BCSCs tumors which is further supported by previous literature (37).

CSCs reside in a special microenvironmental niche that provides the major cues for promoting survival and maintenance (26, 38). The expression of hyaluronan synthase 2 (HAS2) involved in production of hyaluronan and collagen (COL6A3), being the primary ECM components of CSC niche, also plays a precarious role in CSC enrichment (39). In our study, these genes were found to be highly up-regulated in Hi-BCSC tumors as compared to Lo-BCSC tumors, supporting the evidence that ECM integrity determines the BCSC expansion. In addition, matricellular proteins such as Lumican (LUM), osteonectin (SPARC), osteopontin (SPP1), tenascin (TNC), periostin (POSTN) associated with aggressive behavior of disease, known to induce and form of pre-metastatic niche (4, 40) were also found to be up-regulated in Hi-BCSC tumors, thus suggesting association of matricellular protein expression with BCSC expansion.

Thus, in summary, our data from this study clearly shows that up-regulated expression of genes for growth factors, inflammatory cytokines, hypoxia related factors, epithelial to mesenchymal transition and matricellular proteins in stromal and cancer cells as a part of CSC niche in breast tumors is strongly associated with expanded population of cancer stem cells in these tumors. This imbalanced state of gene expression is possibly the result of continuous signals from stromal cells and cancer cells, which results in the generation/expansion of CSCs or acquisition of CSC like phenotype by other cells in the aggressive type of breast cancer. Accordingly, we propose a model of BCSC origin/expansion during advanced stage of breast cancer (Fig. 5C).

## Conclusions

The string analysis of the gene expression data (Figures 5A and 5B) in our study reveals that the stroma around cancer stem cells release many factors that promote the growth of BCSCs while the BCSCs might control metastasis by inducing stemness in the surrounding cancer cells/normal cells by controlling the tumor microenvironment. Accordingly, we may propose that the growth factors along with inflammatory cytokines through Notch and SMAD signaling induce EMT-transcription factors which in turn initiate epithelial to mesenchymal transition to help in enrichment of CSCs.

## Abbreviations

APC-H7: Allophycocyanin-H7
BCSC(s): Breast Cancer Stem Cell(s)
Bp: Base Pair
BV-421: Brilliant Violet-421
CAF(s): Cancer Associated Fibroblast(s)
CD: Cluster of Differentiation
cDNA: Complementary Deoxyribonucleic Acid
CSC(s): Cancer Stem Cell(s)
Ct: Threshold Cycle
CXCL: Chemokine (C-X-C) Ligand
CXCR: Chemokine (C-X-C) Receptor
CXCR4: Chemokine (C-X-C) Receptor 4
DCIS: Ductal Carcinoma *In Situ*
DMEM: Dulbecco’s Modified Eagle Medium
DNA: Deoxyribonucleic Acid
EMP: Epithelial Mesenchymal Plasticity
EMT: Epithelial Mesenchymal Transition
FACS: Fluorescence-Activated Cell Sorting
FBS: Fetal Bovine Serum
FITC: Fluorescein Isothiocyanate
FMO: Fluorescence Minus One
FSC: Forward Scatter
Hi-BCSCs: Tumors with >5% BCSCs
Hi-BCSCs_CC: Cancer Cells isolated from Hi-BCSCs Tumors
Hi-BCSCs_SC: Stromal Cells isolated from Hi-BCSCs Tumors
HIF(s): Hypoxia Inducible Factor(s)
IC-SCR: Institutional Committee on Stem Cell Research
IDC: Invasive Ductal Carcinoma
IEC: Institutional Ethics Committee
IL: Interleukin
LAC: Lumpectomy with Axillary Clearance
LN(s): Lymph Node(s)
Lo-BCSCs: Tumors with <5% BCSCs
Lo-BCSCs_CC: Cancer Cells isolated from Lo-BCSCs Tumors
Lo-BCSCs_SC: Stromal Cells isolated from Lo-BCSCs Tumors
MFI: Mean Fluorescence Intensity
MMP: Matrix Metalloproteinase
MPL(s): Multiple Primary Lesion(s)
mRNA: Messenger Ribonucleic Acid
MSC(s): Mesenchymal Stem Cell(s)
OPD: Out-Patient Department
PCR: Polymerase Chain Reaction
PE: Phycoerythrin
qPCR: Quantitative PCR
RNA: Ribonucleic Acid
RT-PCR: Reverse Transcriptase-Polymerase Chain Reaction
SDF-1: Stromal Derived Factor-1
SSC: Side Scatter
TMAC: Total Mastectomy and Axillary Clearance
TNBC: Triple Negative Breast Cancer
TRI: Trizol Reagent

## Declarations

## Acknowledgments

The authors gratefully acknowledge the help by Dr Meenakshi Sachdeva, DHR Women Scientist in Department of Translational & Regenerative Medicine, PGIMER, Chandigarh for compiling the figures as per journal style and reading the manuscript. The authors are thankful to the technical staff of Histopathology department for help in immunohistochemistry staining.

## Ethics approval and consent to participate

The study was reviewed and approved by Institutes Ethics Committee (ref. no. PGI/IEC/2014/2151 dated 06.01.2014) and Institutional committee on Stem cell Research (IC-SCR) (ref no. PGI-IC-SCRT-56-2013/4516 dated 04.11.2013), Postgraduate Institute of Medical Education and Research, Chandigarh, India. All samples from the patients were taken after informed consent and ethical permission was obtained for participation in the study.

## Consent for publication

An informed consent was obtained from all study subjects for confidentiality of data sharing. The presented data are anonymized and risk of identification is low.

## Availability of data and materials

Technical appendix, statistical code and dataset are available from corresponding author at.

## Authors Contribution

SKA Conceived and designed the experiments; ND: Performed the experiments & data acquisition; ND, SKA, AB: Data analysis and validation. SKA, GS: reagents/patient material/analysis tools; ND, AB, GS, SKA: Drafted the article and made critical revisions related to the intellectual content of the manuscript and approved the final version of the article to be published. All authors meet the three authorship requirements for this journal.

## Captions for supporting information files

Supplementary Table 1: List of selected genes based on their involvement in expansion of CSCs for custom PCR array

Supplementary Table 2: Gene symbols and their official full names.

Supplementary Table 3: Custom PCR Array plate design and format.

Supplementary Table 4: Percentage of BCSCs identified by flow cytometry and IHC in tumor and adjacent normal tissue

